# Identification of genome-wide distribution of cyanobacterial group 2 sigma factor SigE accountable for its regulon

**DOI:** 10.1101/2021.12.09.472044

**Authors:** Ryo Kariyazono, Takashi Osanai

## Abstract

Sigma factors are the subunits of bacterial RNA polymerase that govern the expression of genes by recognizing the promoter sequence. Cyanobacteria, which are oxygenic phototrophic eubacteria, have multiple alternative sigma factors that respond to various environmental stresses. The subgroup highly homologous to the primary sigma factor (SigA) is called the group 2 sigma factor. The model cyanobacterium, *Synechocystis* sp. PCC 6803, has four group 2 sigma factors (SigB-E) conserved within the phylum Cyanobacteria. Among the group 2 sigma factors in *Synechocystis* sp. PCC 6803, SigE is unique because it alters metabolism by inducing the expression of genes related to sugar catabolism and nitrogen metabolism. However, the features of promoter sequence of the SigE regulon remains elusive. Here, we identified the direct targets of SigA and SigE by chromatin immunoprecipitation sequencing (ChIP-seq). We then showed that the binding sites of SigE and SigA overlapped substantially, but SigE exclusively localized to SigE-dependent promoters. We also found consensus sequences from SigE-dependent promoters and confirmed their importance. ChIP-seq analysis showed both the redundancy and specificity of SigE compared with SigA, integrating information obtained from a previously adopted genetic approach and *in vitro* assays. The features of SigE elucidated in our study indicate its similarity with group 2 sigma factors of other bacteria, even though they are evolutionally irrelevant. Our approach is also applicable to other organisms and organelles, such as plant plastids, which have multiple group 2 sigma factors.

**Importance:** Group 2 sigma factors are alternative sigma factors highly homologous to primary sigma factors. Cyanobacteria, which are photosynthetic eubacteria, are unique because they have multiple group 2 sigma factors. Although each sigma factor induces the expression of specific genes, the redundancy and complicated network of the primary and group 2 sigma factors hinder the identification of their regulons *via* a genetic approach. Here, we identified the binding sites of SigE (group 2 sigma factor) and SigA (primary sigma factor) using chromatin immunoprecipitation sequencing and identified the minimal element of SigE-dependent promoters by subsequent promoter assays. Our study provides insights into the common features of group 2 sigma factors, which, though evolutionarily irrelevant, are widespread among eubacteria and plant plastids.

## Introduction

Cyanobacteria are oxygenic phototrophic eubacteria that inhabit a wide range of environments. Similar to other organisms, cyanobacteria respond to environmental changes by altering their gene expression profile. The protein complex RNA polymerase (RNAP), which performs transcription, comprises the core enzyme and the dissociable subunit sigma factor (1). Sigma factors determine the genes to be transcribed by preferentially binding to promoters in cooperation with transcription factors (2). The promoters of typical housekeeping genes possess consensus sequences at 10 and 35 bp upstream of the transcription start sites (TSS). Comparative evaluation of the active promoters of *Escherichia coli* genes helped identify -10 consensus (TATAAT) and -35 consensus (TTGACA) sequences with some deviation (3). These consensus sequences also exist in a subset of cyanobacterial promoters (4, 5), and heterologous promoters act reciprocally in cyanobacteria and *E. coli* (6, 7), implying that the promoter consensus sequences are valid for cyanobacteria.

Usually, bacterial species possess the primary sigma factor responsible for the transcription of housekeeping genes and several alternative sigma factors that respond to environmental stresses (8, 9). Some alternative sigma factors have a consensus sequence distinct from that of the primary sigma factor, whereas others have a consensus sequence that is only slightly different from that of the primary sigma factor. While the primary sigma factor is well conserved among eubacteria, alternative sigma factors are unique to clades and vary in their number among species. The phylum Cyanobacteria possess one primary sigma factor (group 1), several sigma factors highly homologous to the primary sigma factor (group 2), sigma factors with limited homology to the primary sigma factor (group 3), and extracytoplasmic function sigma factors (group 4) (10, 11).

*Synechocystis* sp. PCC 6803 (hereafter referred to as *Synechocystis*) is a unicellular freshwater cyanobacterium that was the first cyanobacterium to undergo complete genome sequencing (12); hence, investigations on its sigma factors have markedly advanced. *Synechocystis* produces nine sigma factors: one primary sigma factor (SigA), four group 2 sigma factors (SigB, C, D, and E), one group 3 sigma factor (SigF), and three group 4 sigma factors (SigG, H, and I). Except for SigG, alternative sigma factors of *Synechocystis* have been experimentally proven to be non-essential (13, 14). Each group 2 sigma factor responds to a different spectrum of environmental stress, but some stress conditions induce responses from multiple sigma factors (14–17). Although group 2 sigma factors respond to stress conditions, they are incorporated into the RNAP complex and participate in transcription under optimal growth conditions (18).

Cyanobacterial promoters are classified into three types (19): type 1 promoters equipped with -10 and -35 consensus sequences, type 2 promoters with a -10 consensus sequence but no canonical -35 consensus sequence, and type 3 promoters with distinct consensus sequences specifically recognized by SigF (20). Some type 2 promoters have binding sites for transcription factors adjacent to the -35 region instead of the consensus sequence (19). SigA and group 2 sigma factors recognize both type 1 and type 2 promoters, but group 2 sigma factors seem suitable for inducing gene expression *via* type 2 promoters (21, 22). For example, SigD can recognize the *psbA2* promoter (type 1) independent of the -35 consensus sequence (23), and SigB and SigC recognize type 2 promoters in cooperation with the transcription factor NtcA that binds to a site adjacent to the -35 region (17, 24). Owing to the variable -35 region of type 2 promoters and redundant recognition by sigma factors, the consensus sequence associated with the specific regulon of group 2 sigma factors remains elusive.

In addition to the redundant regulon, the competition and hierarchy of sigma factors complicate their relationship. Sigma factors compete for the core enzyme of RNAP, enabling acute regulation in response to environmental changes (8, 22, 25). Some sigma factors are responsible for the expression of other sigma factors (24). This complicated network of sigma factor expression makes it difficult to identify the specific regulon of group 2 sigma factors *via* a genetic approach. The binding sites of sigma factors can be directly recognized by chromatin immunoprecipitation (ChIP) sequencing (ChIP-seq) combined with the conventional genetic approach.

Among alternative sigma factors of *Synechocystis*, SigE is unique because it alters carbon metabolism by inducing the expression of genes related to glycolysis, the oxidative pentose phosphate (OPP) pathway, and glycogen catabolism (26). Although SigE shows activity under continuous light exposure, SigE gene and protein expression is regulated by the circadian rhythm (27), and light exposure suppresses SigE protein activity (28). Through these regulations, SigE activates sugar catabolism-related genes when photosynthetic growth is disabled. Additionally, *sigE* expression is positively regulated by the global nitrogen regulator NtcA (17). This regulation may help maintain the carbon-nitrogen balance by altering metabolism under conditions of nitrogen starvation (29, 30). SigE is widely distributed among cyanobacteria, except in the subgroups of marine cyanobacteria, which implies that SigE plays a pivotal role in the cyanobacterial clade (19).

The consensus sequence of SigE-driven promoters remains elusive because *in vitro* transcription assays have shown that primary and group 2 sigma factors recognize promoters redundantly, whereas the genetic approach has revealed that SigE recognizes the promoters of specific genes; however, SigE induces a distinct expression profile. To determine the promoter regions that SigE recognizes, we performed ChIP-seq analysis and identified the SigE-binding region on a genome-wide scale. We found that SigE could bind to the promoter region of both SigE-dependent and -independent genes, and most binding sites of SigE overlapped with those of SigA, but SigE specifically binds to SigE-dependent genes. We propose that the expression of SigE regulons is the outcome of the balance between the affinities of SigE and other sigma factors to the promoter and determined by its sequence.

## Results

### Majority of SigE- and SigA-binding sites overlap

To determine the genome-wide distribution of SigE- and SigA-binding sites, we conducted ChIP-seq of FLAG-tagged SigE and SigA strains (Fig. S1 and S2). We identified 478 SigE-binding sites and 859 SigA-binding sites from two biological replicates and one un-tagged control immunoprecipitation (IP) (Fig. S2C). Of the 478 SigE sites, 394 overlapped with SigA sites (Fig. 1A), and the strong peaks tended to overlap (Fig. 1B and C Next, we compared the positions of previously identified TSS (31) and SigA- or SigE-binding sites and found that the majority of the binding summits of SigE and SigA (90.0% and 87.3%, respectively) were located within 100 bp upstream or downstream of the TSS (Fig. S3A).

**Figure 1.**
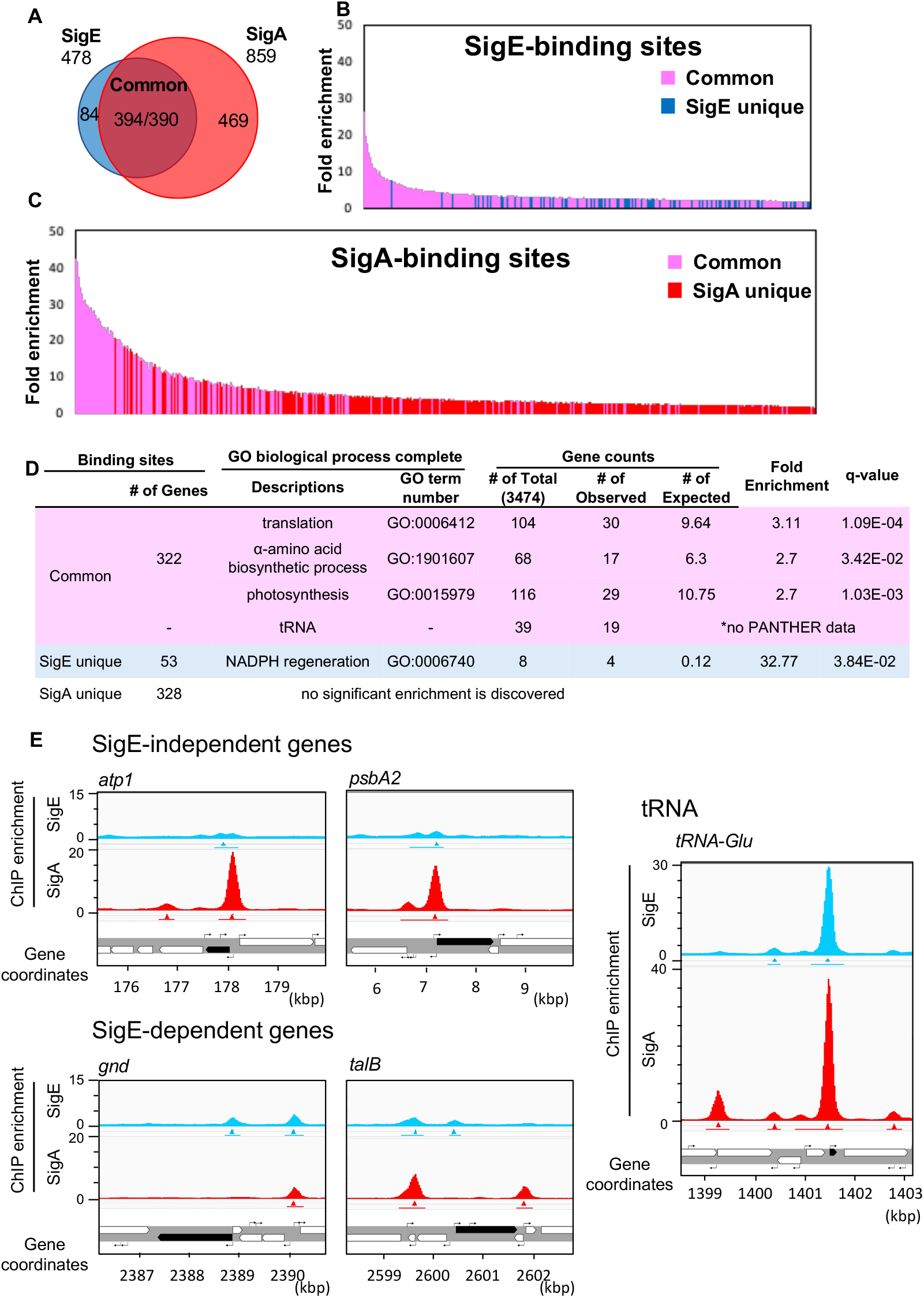
Genome-wide occupancy of SigA and SigE analyzed using chromatin immunoprecipitation sequencing (ChIP-seq) analysis. (A) Venn diagram showing the overlap of SigA- and SigE-binding sites. “394/390” in the region of intersection in the Venn diagram indicates the overlap between 394 SigE-binding sites and 390 SigA-binding sites. (B) Binding sites of SigE in the order of immunoprecipitation (IP) enrichment. The regions overlapping with SigA-binding sites are highlighted in magenta, and the regions devoid of SigA-binding sites are highlighted in blue. (C) Binding sites of SigA in the order of IP enrichment. The regions overlapping with SigE-binding sites are highlighted in magenta, and the regions devoid of SigE-binding sites are highlighted in red. (D) Results of gene ontology (GO) enrichment analysis for genes downstream of common binding sites for SigE and SigA, SigE unique sites, and SigA unique sites. Only GOs from the bottom of the hierarchy are listed. The “q-value” indicates the false discovery ratio. (E) ChIP enrichment of SigA and SigE in the regions surrounding the SigE-independent genes (top left; *atp1* and *psbA2*), SigE-dependent genes (bottom left; *gnd* and *talB*), and tRNA-coding genes (right). ChIP fold enrichment of SigE (top) and SigA (middle) and the gene coordinates (bottom) are shown. The peak regions and summits are indicated as bars and triangles below the fold enrichment graph. The black arrows in the gene coordinate row indicate the genes of interest. The hooked arrow indicates the transcription start site.

### Housekeeping genes among genes downstream of common binding sites are enriched

As most of the binding sites of SigE and SigA are located close to TSS, we assigned each binding site to the transcription units (TUs) and attempted to identify the genes downstream of the binding sites. We applied gene ontology (GO) enrichment analysis (32) against three sets of genes: genes downstream of sites corresponding to SigE unique peaks, peaks common to SigE and SigA, and SigA unique peaks. We found that genes related to translation (GO:0006412), α-amino acid biosynthetic process (GO:1901607), and photosynthesis (GO:0015979) were enriched downstream of the sites associated with the common peaks, and genes associated with NADPH regeneration (GO:0006740) were enriched downstream of the sites associated with the SigE unique peaks, while no significant enrichment was observed downstream of the sites associated with the SigA unique peaks (Fig. 1D). In addition to protein-coding genes, genes coding tRNA and rRNA were preferentially detected in common peaks (19 of 39 tRNA genes and both 16S-tRNA-23S-5S rRNA clusters). The enriched genes in common peaks tended to show a high binding score for both SigE and SigA (Fig. S3B).

### SigE specifically recognizes SigE-dependent promoters

GO enrichment analysis showed that four genes related to NADPH regeneration, namely, *talB* (slr1793), *gnd* (sll0329), *pntA* (slr1239), and *zwf* (slr1843), lie within SigE unique binding sites, which are known to be regulated by SigE. Therefore, we speculated that genes transcribed in a SigE-dependent manner prefer SigE, but not SigA (Fig. 1E). To confirm this, we listed genes identified in previous studies that reportedly undergo SigE-dependent transcription (26, 30), i.e., genes whose expression is reduced in response to *sigE* deletion (<-0.5 in log_2_ scale fold change) and enhanced in response to SigE overexpression (>0.5 in log_2_ scale fold change). We referred to these as SigE-dependent genes and listed the corresponding TUs from a previous study (31) (listed in Table 1).

**Table 1.**
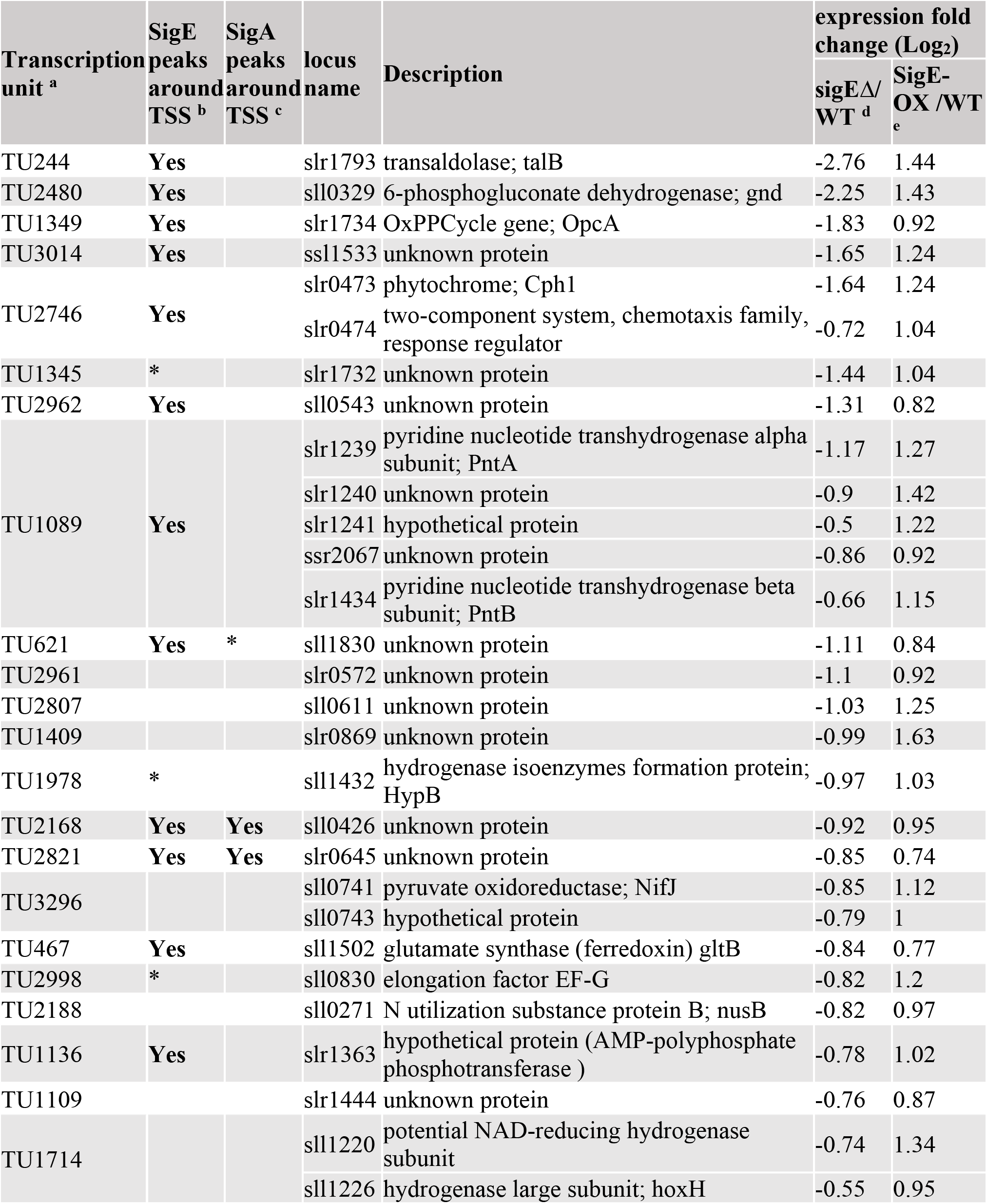
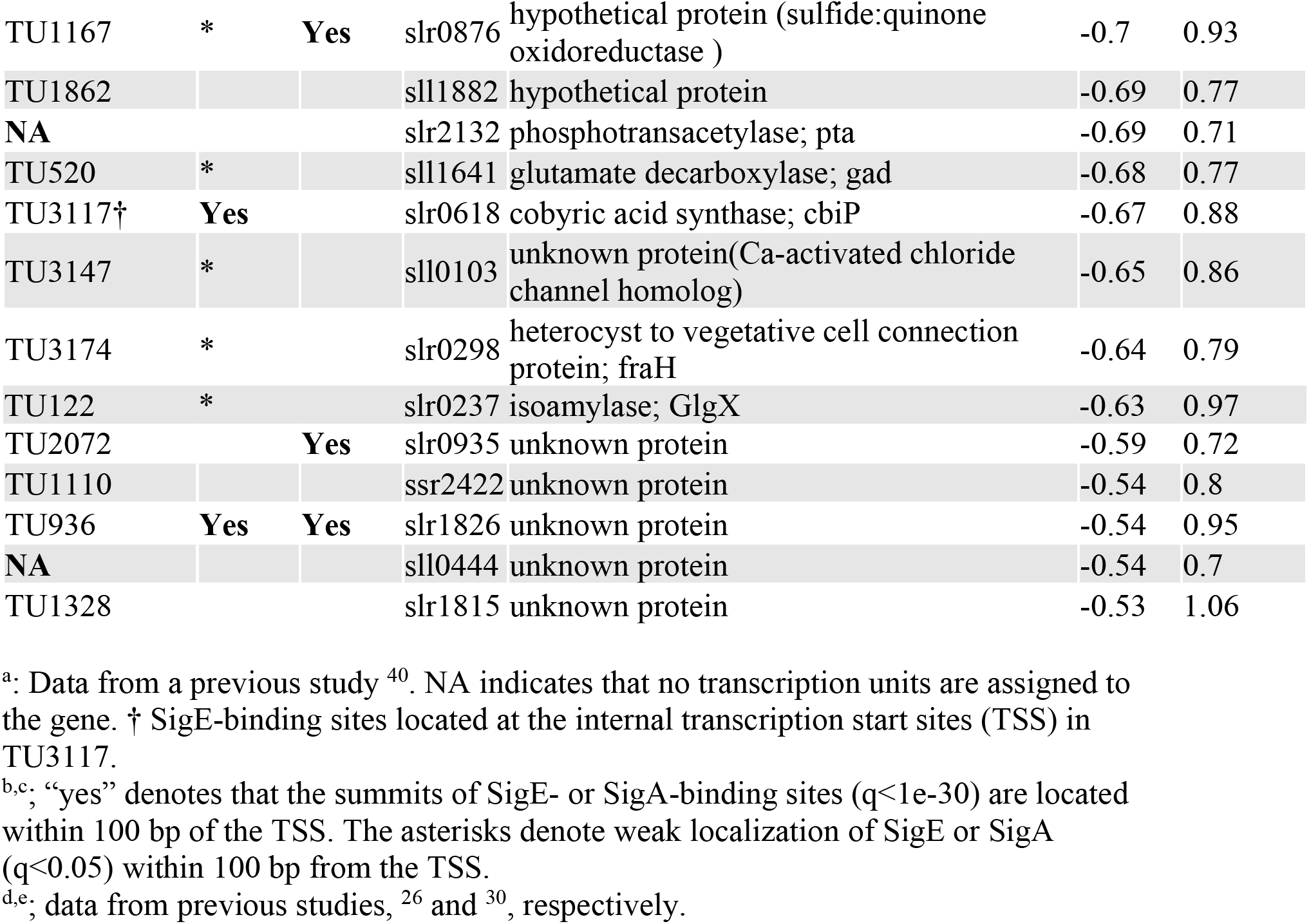
List of SigE-dependent genes. The selected genes are sorted by the transcription units and in the order of fold enrichment of gene expression in *sigEΔ* cells.

Contrary to the general similarities between SigE- and SigA-binding sites, the TSS of SigE-dependent genes were enriched in SigE-binding sites but not in SigA-binding sites; 14 of 33 TSS belonged to SigE-binding sites, whereas six TSS belonged to SigA-binding sites (Fig. S3B and Table 1). ChIP enrichment of SigE-binding sites assigned to SigE-dependent genes tended to be weaker than those assigned to genes encoding tRNA, rRNA, and housekeeping genes (Fig. S3B). From the analysis, we concluded that SigE, rather than SigA, bound to the promoters of SigE-dependent genes, although we noted that 8 of 33 TSS in SigE-dependent genes showed weak localization of SigE (peaks detected under the threshold of q<0.05), whereas 11 of 33 TSS did not show significant accumulation of SigE (Table 1).

### Motif identification in SigE-dependent promoters

To date, we have elucidated that most of the binding sites of SigE and SigA are common, but the binding preferences are slightly different, which leads to SigE dependence. To understand this difference, we searched the consensus motif around SigA- and SigE-binding summits using the STREME program (33). We identified -10 consensus sequences and extended motif (a<u>**tG**</u>N<u>**TA**</u>NNN<u>**T**</u>) distributed around the center of SigA peaks, and similar motifs were found for SigE (<u>**G**</u>N<u>**T**</u>ANNA<u>**T**</u>GG) (Fig. 2A). When we focused on only strong SigA-binding sites (>20 IP fold enrichment), another consensus sequence (TTNG/A) was identified at the -35 position (Fig. 2B), which partially matched the canonical -35 consensus sequence in *E. coli* (<u>**TT**</u>G<u>**A**</u>CA).

**Figure 2.**
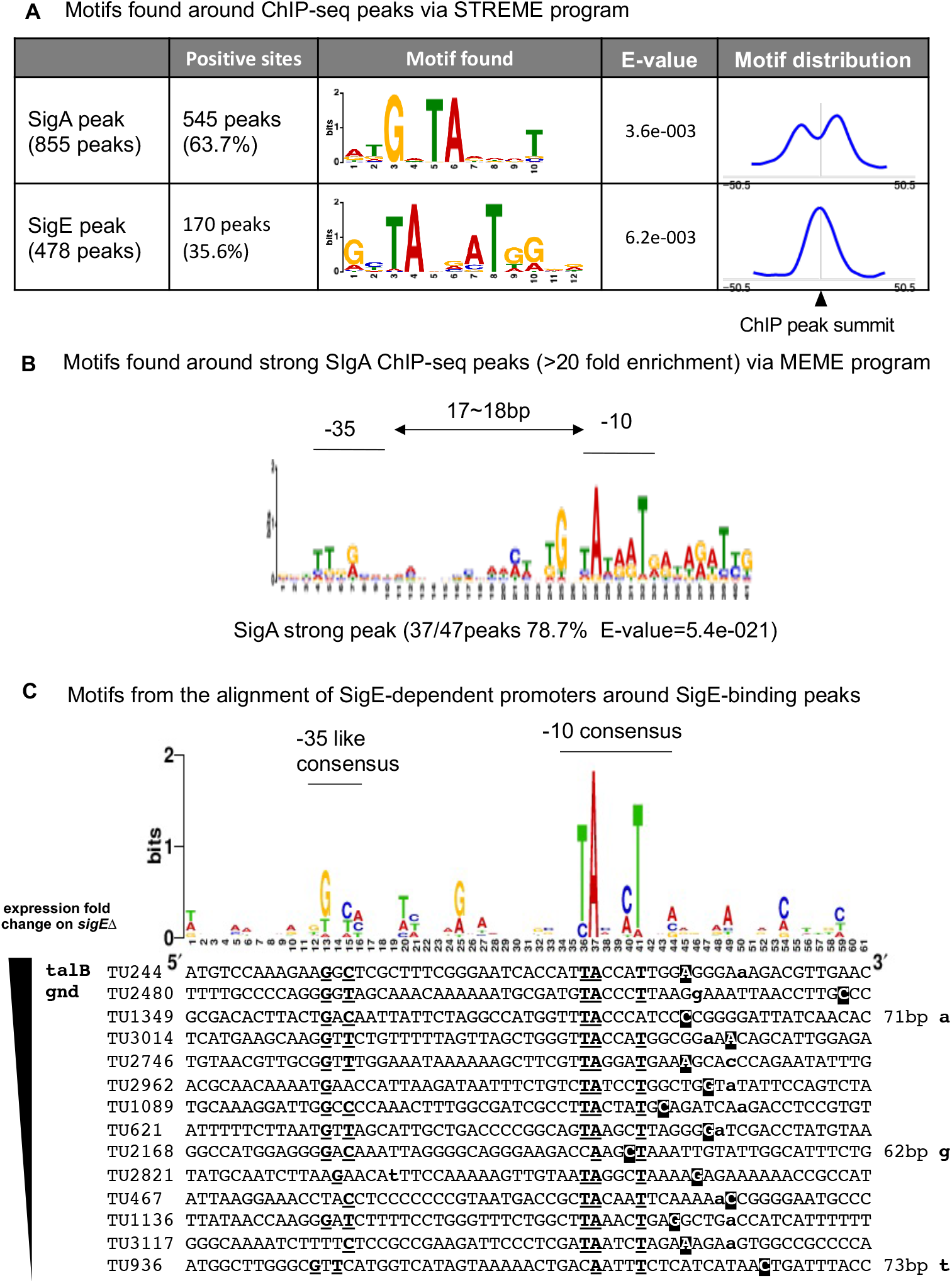
(A) Sequence logo from sequences around all SigA-binding sites (top) and SigE-binding sites (bottom). The motifs were identified using the STREME program. Motif distribution is shown in the right column. (B) Sequence logo from strong SigA-binding sites (>20 IP fold enrichment). The motifs were identified using the MEME program. (C) Sequence logo from 14 SigE-dependent promoters with SigE-binding sites (top); the alignment of 14 SigE-dependent promoters (bottom). The sequences are ordered with respect to the fold change of expression in *sigE*Δ. The positions of transcription start sites are indicated by boldfaced lowercase letters. The SigE-binding summits are indicated by white letters against a black background. The consensus nucleotides are indicated by boldfaced underlined letters.

Next, we searched for the consensus sequence of the SigE regulon. To this end, we selected SigE-binding sites near the TSS of SigE-dependent genes. From the alignment of 14 sequences, we identified a consensus sequence at positions corresponding to the -10 and -35 sites. The consensus sequence at the -10 site showed a degenerated canonical -10 consensus sequence (TANNNT) with cytosine-enriched NNN, and an extended -10 consensus sequence did not appear. Moreover, in the region corresponding to the -35 site, we identified a consensus sequence (GNC/TNNN) considerably different from the canonical -35 consensus sequence (TTGACA); we hereafter refer to it as “-35G” (Fig. 2C).

### Promoter assay confirmed the essential element in SigE-dependent promoters

From the alignment of the SigE regulon, we estimated the positions of the -10 consensus sequence and -35G of SigE-dependent promoters (Fig. 2C and 3A). To confirm that the estimated consensus sequence is important for SigE-dependent promoters, we conducted a promoter assay by measuring the green fluorescent protein (GFP) content in the presence or absence of SigE. We cloned the upstream region of *gnd* (sll0329) and *talB* (slr1793) as typical SigE-dependent promoters (*Pgnd* and *PtalB*, respectively) (Fig. 3A) and confirmed that *Pgnd* and *PtalB* behave in a SigE-dependent manner in this assay (Fig. 3B and 3C). We then demonstrated that mutations in the predicted -10 consensus sequence induced GFP expression at undetectable levels, and mutations in -35G suppressed promoter activity (Fig. 3B and 3C). Next, we exchanged the -10 consensus sequence of *Pgnd* with that of housekeeping promoter, *Patp1* (TA<u>CCC</u>T to TA<u>TGA</u>T).

**Figure 3.**
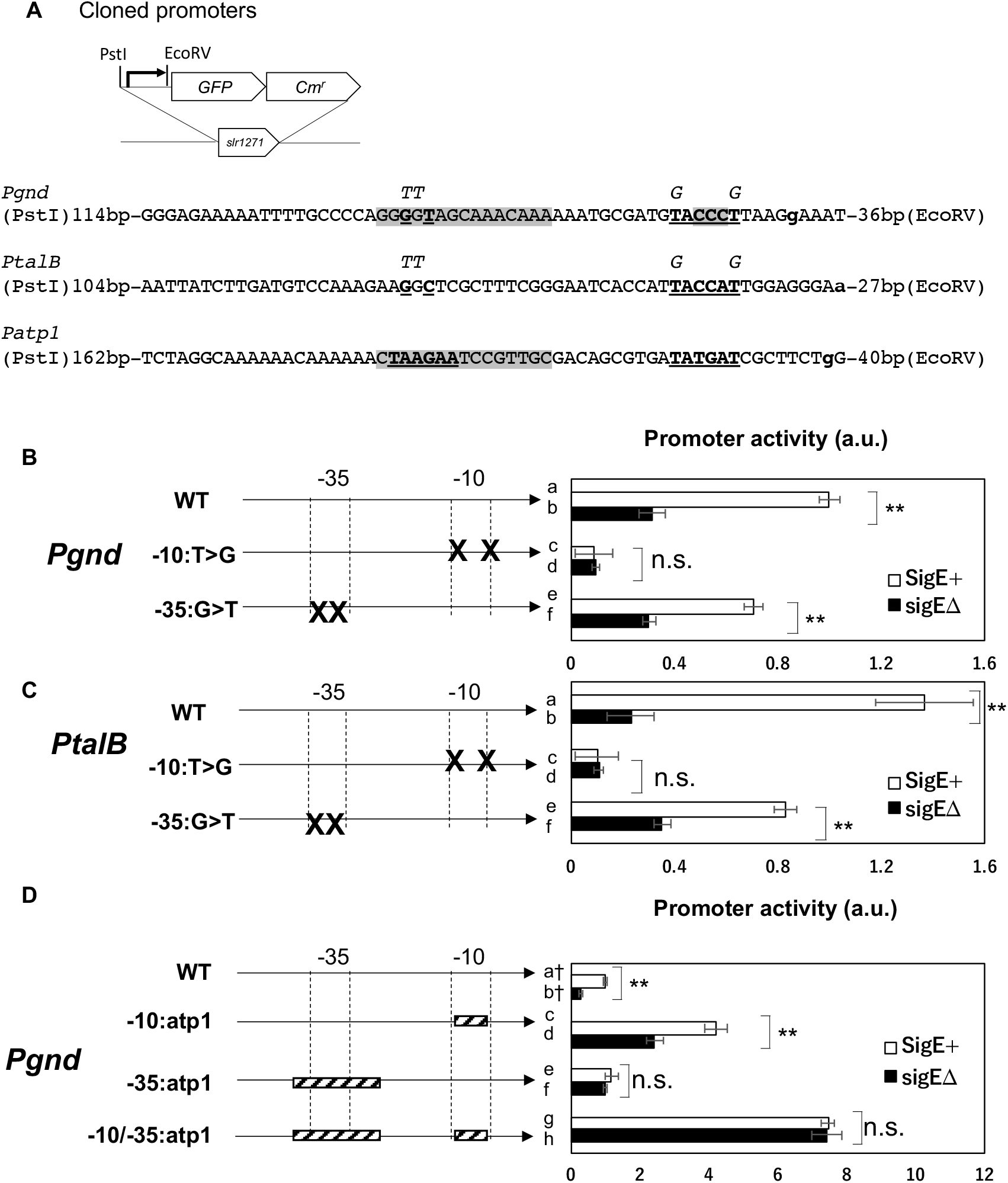
The activity of promoters in the presence or absence of SigE. Promoter activity was measured based on the quantity of GFP detected using western blotting. The values are normalized with respect to those of *Pgnd*-GFP in the SigE+ strain (KR5). Error bars indicate the standard deviations from values obtained from three biological replicates. The asterisks denote the statistical significance between the values obtained for SigE+ and *sigE*Δ strains determined by pairwise comparison via the Student’s *t*-test (* p<0.05, ** p<0.01, n.s. not significant.) The schematic representation of the positions of mutation(s) or replacement(s) is provided on the left. (A) Schematic representation of the promoter-GFP construct integrated into the cyanobacterial genome, and the sequence of promoters analyzed in this assay. The positions of transcription start sites are marked using boldfaced lowercase letters, and the -10 and -35 consensus sequences are marked using boldfaced underlined letters. The italicized letters on the sequence indicate mutations introduced in (B) and (C). The shaded regions show 16 nucleotides replaced with corresponding sequences from *Patp1* in (D). (B) The activity of the *gnd* promoter and their mutant derivatives in the presence or absence of SigE. The letter “X” in the left diagram indicates the mutated nucleotides. The strains shown in this figure are: a. KR5, b. KR6, c. KR16, d. KR17, e. KR193, and f. KR194 (C) The activity of the *talB* promoter and their mutant derivatives in the presence or absence of SigE. The letter “X” in the left diagram indicates the mutated nucleotides. The strains shown in this figure are: a. KR30, b. KR31, c. KR161, d KR162, e. KR241, and f. KR242 (D) The activity of the *gnd* promoter and its derivatives, whose consensus sequence was replaced with that of *Patp1*. The striped boxes in the left diagram indicate the region replaced with the corresponding region of *Patp1*. The strains shown in this figure are: a. KR5, b. KR6, c. KR139, d. KR140, e. KR249, f. KR250, g. KR251, and h. KR252. †: The values of KR5 and KR6 are the same as those in (B).

This modification promoted activity both in the presence and absence of SigE (Fig. 3D c and d). The replacement of the 16 nt region, which included -35G, led to the loss of SigE specificity (Fig. 3D e and f) but did not enhance promoter activity. The combined replacement of the -10 consensus sequence and 16 nt region around -35G further enhanced promoter activity and led to the loss of SigE specificity. The findings associated with the mutation and replacement of *Pgnd* indicate that the -10 consensus sequence prevents SigE-independent high gene expression. Furthermore, the 16 nt region containing -35G, rather than -35G alone, reinforces SigE specificity.

### Minimal sufficient element of SigE-dependent promoters

To determine the minimal sufficient element of SigE-dependent promoters, we modified the sequence of SigE-independent promoters to induce SigE-dependent behavior. We cloned the upstream region of *atp1* (sll1321) and *psbA2* (slr1311) for the SigE-independent promoter (*Patp1* and *PpsbA2*, respectively, Fig. 4A) and confirmed that *Patp1* and *PpsbA2* expressed GFP both in the presence and absence of SigE under continuous light exposure (Fig. 4B and C). The activity of *Patp1* and *PpsbA2* was considerably higher than that of SigE-dependent promoters, consistent with the levels of the transcripts produced (31, 34).

**Figure 4.**
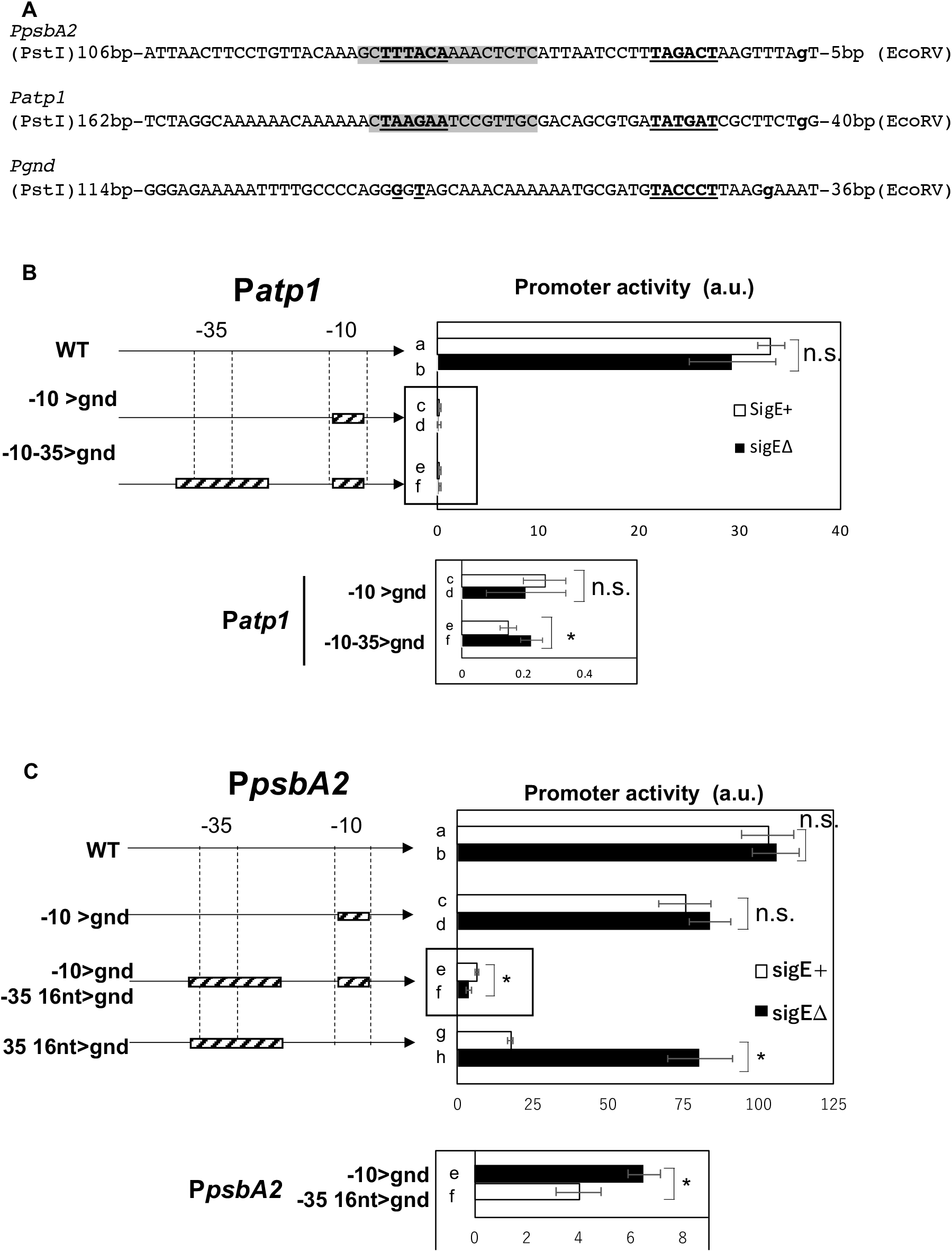
The activity of the promoters indicated in Figure 3. The values are normalized to those obtained for *Pgnd*-GFP in the SigE+ strain (KR5), as shown in Figure 3B. (A) Sequence of the promoters analyzed in this assay. The positions of the transcription start sites are marked using boldfaced lowercase letters, and the -10 and -35 consensus sequences are marked using boldfaced underlined letters. The shaded regions show the 16 nucleotides replaced with the corresponding nucleotides of *Pgnd*. (B) The activity of the *atp1* promoter and its derivatives, whose consensus sequences were replaced with those of *Pgnd*. The striped boxes in the left diagram indicate the region replaced with the corresponding region of *Pgnd*. The strains assayed were: a. KR127, b. KR128, c. KR145, d. KR146, e. KR189, and f. KR190. The box below the graph shows the magnified graph of mutant promoters. (C) The activity of the *psbA2* promoter and its derivatives, whose consensus sequence was replaced with that of *Pgnd*. Striped boxes in the left diagram indicate the region replaced with the corresponding region of *Pgnd*. The strains assayed were: a. KR9, b. KR10, c. KR24, d. KR25, e. KR78, f. KR79, g. KR56, and h. KR57. The box below the graph shows the magnified graph of e.KR78 and f. KR79.

When we replaced the -10 consensus sequence of *Patp1* with that of *Pgnd*, promoter activity decreased to an undetectable level (Fig. 4B c and d). In addition to replacing the - 10 consensus sequence, we further replaced the 16 nt region of P*atp1* around the -35 consensus sequence with that of *Pgnd*, but the modified promoter did not show SigE dependence (Fig. 4B e and f). In case of *PpsbA2*, the replacement of the -10 consensus sequence retained expression both in the presence and absence of SigE (Fig. 4C c and d), and the simultaneous replacement of the 16 nt region around the -35 consensus sequence was sufficient to induce SigE-dependent behavior in the promoter (Fig. 4C g and h), although this promoter remained active in absence of SigE. We succeeded in rendering SigE dependence in one promoter but failed in the case of the other promoter.

### Roles of different elements in SigE-dependent promoters

We concluded that the -10 region is the primary determinant of SigE specificity owing to its ability to limit access to SigA (Fig. 5) for the following reasons: 1) the more stringent SigE-dependent promoters are, the greater is the cytosine enrichment in the -10 consensus sequence (Fig. 2C); 2) replacement of the -10 consensus region in *Pgnd* with that of *Patp1* increased promoter activity and reduced SigE specificity (Fig. 3D and 5A); and 3) replacement of the -10 consensus sequence of *Patp1* with that of *Pgnd* markedly decreased promoter activity (Fig. 4B and 5B). *PpsbA2* retained its promoter activity when the -10 consensus sequence was replaced with that of *Pgnd* because it may have been recognized by SigD. The region containing the -35 consensus sequence also determines SigE specificity and supports SigE-dependent promoter activity (Fig. 2B and 2C). In addition to the -10 and -35 consensus sequences, the other region including upstream sequence is required for the optimal activity of SigE-dependent promoters (Fig. 5C).

**Figure 5.**
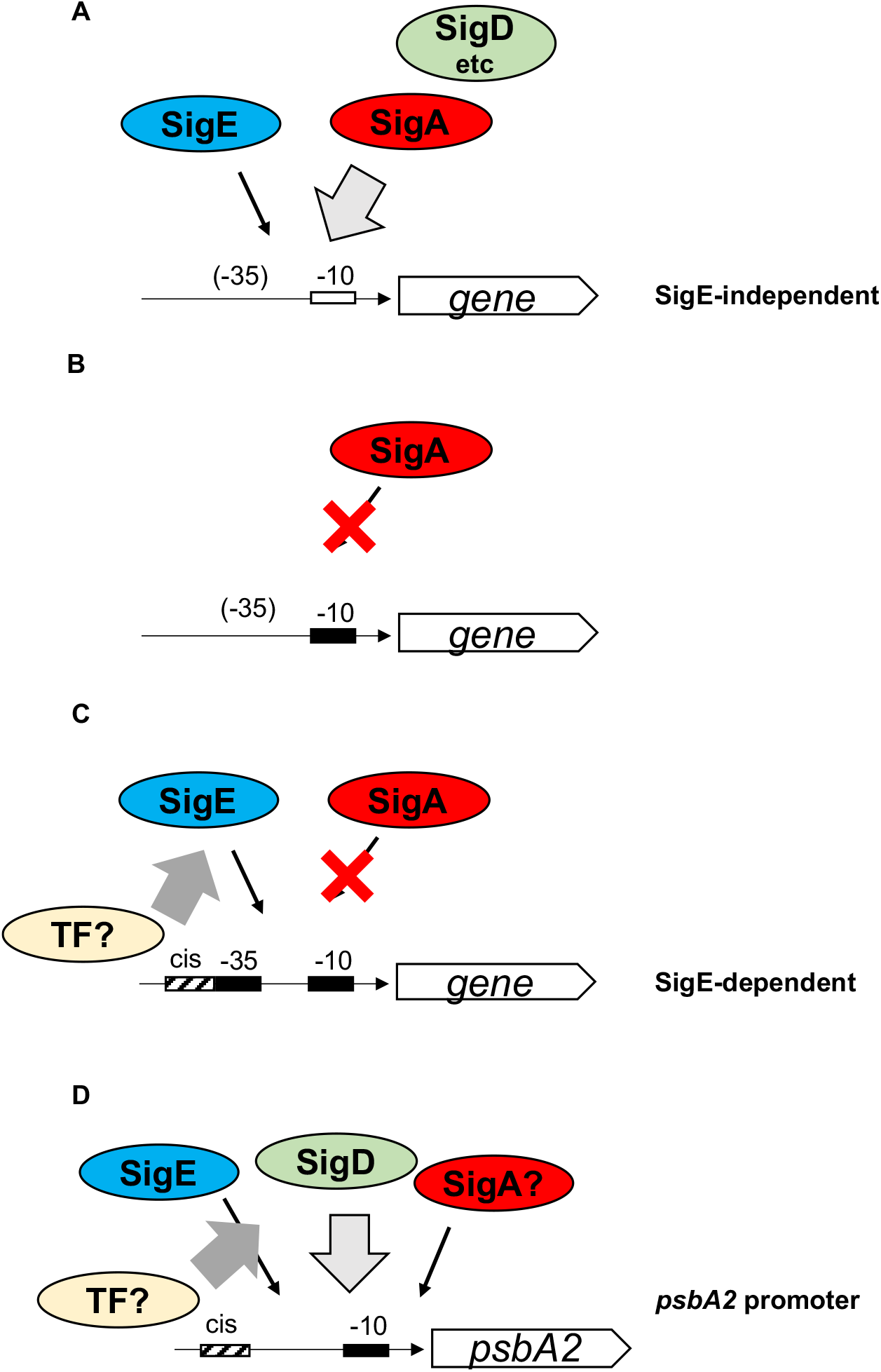
Schematic diagram of the gene promoters recognized by SigE and SigA. (A) Promoter of SigE-independent genes, which is recognized by both SigA and SigE (and presumably by other sigma factors). The white box shows the canonical -10 consensus sequences. (B) SigA cannot recognize degenerated -10 consensus sequences, indicated by the black box. (C) SigE can recognize promoters with the consensus sequences identified in this study, indicated by the black boxes. The cis-element upstream of the -35 consensus sequences indicated by the striped box may recruit transcription factors (denoted as TFs) and support SigE. (D) In the case of the *psbA2* promoter, in which the - 10 consensus sequences were replaced with those of *Pgnd*, the TF may support transcription *via* SigE.

## Discussion

The present study identified the binding sites of the group 2 sigma factor, SigE, and the primary sigma factor, SigA, *via* ChIP-seq analysis. We found that strong binding sites were common for SigE and SigA, but SigE-dependent promoters preferred binding to SigE rather than SigA. We also identified a consensus sequence of SigE-dependent promoters that was experimentally verified to be important for SigE-dependent transcription.

### ChIP-seq analysis as a reliable tool for regulon analysis

To date, two types of conventional approaches have been adopted to evaluate the contribution of sigma factors with overlapping regulons: transcriptome analysis and *in vitro* transcription assays. While transcriptome analysis of sigma-disrupted strains can help detect genes that are highly dependent on each sigma factor *in vivo*, other sigma factors might compensate by inducing the expression of some genes. This genetic approach has helped to successfully identify the consensus sequence of group 3 and 4 sigma factors (20, 35) owing to their high specificity. *In vitro* transcription assays using the reconstituted RNAP core enzyme and sigma factors help assess the promoter selectivity of individual sigma factors; however, it is difficult to take into account the effect from other sigma factors. ChIP-seq analysis can support these conventional assays by specifying the direct target genes for the sigma factor of interest *in vivo*. Our ChIP-seq approach, when assisted by transcriptome analysis of sigma-disrupted or -overexpressing strains, can be applied to other sigma factors. Combination with data on TSS will also strongly support the ChIP-seq analysis of sigma factors.

### Requirement of cis-elements for SigE-dependent promoters is presumed

As group 2 sigma factors have structure similar to that of the primary sigma factor, it is expected that SigE directly recognizes the -35 region of the promoter but not the region upstream of the -35 site. However, our promoter assay showed that when the -10 consensus sequence was replaced, the -35 consensus sequence was insufficient for rendering SigE dependence on the *atp1* promoter (Fig. 4B). This implies that the upstream region of the promoter is necessary and may recruit transcription factors, which cooperate with SigE to transcribe SigE-dependent genes. Conversely, the modified *PpsbA2* showed SigE dependence upon the replacement of the -10 and -35 consensus sequences but not the upstream region of the -35 site (Fig. 4C). The *psbA2* promoter possesses a cis-element upstream of the -35 site (7, 36), which might assist SigE activity at the modified *psbA2* promoter. Although no transcription factor is known to act with SigE, the other group 2 sigma factors are known to cooperate with different transcription factors. For example, SigC cooperates with the transcription factor NtcA and induces transcription from the *glnB* promoter, in which the NtcA-binding site is positioned adjacent to the -35 region (37).

Among the 33 SigE-dependent TUs, eight showed weak SigE localization, and 11 showed no SigE peaks (Table 1). SigE might indirectly promote those TUs; alternatively, SigE recognizes such TUs in cooperation with the transcription factors, which were inactive under the culture conditions in the present ChIP experiment. For example, TU1714, which encodes hydrogenase gene cluster, is devoid of SigE and SigA binding peaks and is expressed under the anaerobic condition with the help of the transcription factor (38).

SigE is regulated by the light-dark cycle and is essential for survival in the dark. In the initial period of darkness, cyanobacteria adapt to the dark; later, respiration makes the environment anaerobic, and as the period of light exposure (i.e., dawn) approaches, the activation of the OPP pathway is necessary for rebooting photosynthesis (39), whereas, nitrogen starvation may occur regardless of the diurnal cycle. SigE may respond to such multifaceted requirements *via* cooperation with the transcription factors responsible for responding to each condition.

### Common features of group 2 sigma factors in eubacteria

The promoter recognition spectrum of SigE is analogous to that of RpoS, a group 2 sigma factor of *E. coli*. Using a genetic approach (40) and ChIP-seq analysis (41, 42), it was demonstrated that RpoS can recognize both housekeeping promoters and unique promoters. In *Mycobacterium*, the group 2 sigma factor σ^B^ transcribes housekeeping genes in a manner similar to the primary sigma factor σ^A^ (43). Similar behavior has been noted in RpoS, σ^B^, and SigE, even though these sigma factors have evolved independently; hence, this might be a universal feature of group 2 sigma factors. In contrast to RpoS and σ^B^ as the sole group 2 sigma factors, cyanobacteria have multiple group 2 sigma factors. Therefore, group 2 sigma factors in cyanobacteria are likely to have a distinct affinity to promoter sequence, not merely tolerant to deviation from canonical consensus, leading to the motif found in this study.

Thus, it is necessary to determine why cyanobacteria possess multiple group 2 sigma factors. Some genes undergo both conditional and basal transcription; for example, *psbA2* expression is upregulated upon high light in addition to basal transcription (19). The redundancy of sequence recognition among the primary and group 2 sigma factors helps shorten the promoters of such genes owing to the sharing of -10 consensus sequences, which help economize genome space. The sharing of the -10 consensus sequence is also beneficial for the fine-tuning of gene expression through microevolution. For example, the modification of a few nucleotides altered the balance between SigE-dependent and basal transcription in our promoter assay (Fig. 2C and 3C).

Plant plastids have sigma factors analogous to the primary sigma factor and multiple group 2 sigma factors (44), and their regulons overlap in some cases, whereas in other cases, the regulons are specific. Although plastid group 2 sigma factors are evolutionally irrelevant to cyanobacterial group 2 sigma factors, ChIP-seq analysis may help shed light on the regulation of plastid sigma factors, as shown for cyanobacteria.

## Materials and methods

### Bacterial strain culture conditions and plasmids

The glucose-tolerant (GT) strain of *Synechocystis* sp. PCC 6803 (45) and its derivatives were cultivated in BG-11_0_ liquid medium (46) containing 5 mM NH_4_Cl and buffered with 20 mM HEPES-KOH (pH 7.8) under continuous exposure to white light (40 μmol m^-2^s^-1^) and bubbled with air containing 1% CO_2_. The SigE (sll1689)-disrupted strain (G50), constructed in a previous study (26), was used. Strains for the promoter assay and epitope-tagged strains were constructed by homologous double recombination between the genome and plasmids (45), and the transformants were selected by three passages on BG-11 plates containing 20 μg/mL chloramphenicol or 5 μg/mL kanamycin. The upstream sequences of *gnd* (sll0329), *talB* (slr1793), *psbA2* (slr1311), and *atp1* (sll1321) were inserted upstream of the region encoding GFP. The “promoter-GFP” constructs were integrated into the genome of the SigE+ strain (GT) or sigE-disrupted strain (G50).The promoter-GFP fragments were inserted into the slr0271 locus, which was reported as one of the “genomic neutral sites” in a previous study (47). Genetic integration of the epitope tag or promoter-GFP fragments was confirmed by genomic polymerase chain reaction (PCR) (Fig. S1A and S4B). The cyanobacterial strains used in this study are listed in Supplemental Table S1.

The plasmids were synthesized by Eurofins Genomics (Japan), amplified from the GT genome by PCR, or mutagenized by site-directed mutagenesis using the PrimeSTAR® Max DNA Polymerase (Takara bio). The plasmids used in this study are listed in Supplementary Table S3.

### Sample preparation for protein assay

Cells were harvested after cultivation for 24 h in HEPES-buffered BG-11_0_ medium containing 5 mM NH_4_Cl (starting from OD_730_ = 0.4), washed once with PBS, and stored at -80 °C. Cells equivalent to those present in 15 mL of the culture were added to 50 μL of the lysis buffer (PBST supplemented with complete and 1 mM PMSF) and 100 μL of acid-washed glass beads (150-212 μm; G1145; Sigma Aldrich). The cell lysates were bead-beaten using ShakeMaster® NEO (Bio Medical Science, Tokyo, Japan) with eight repetitions of 1 min at 1500 rpm and incubation at 1-min intervals on ice. After the debris was removed by centrifugation (18,000 × *g*, 5 min, 4°C), the supernatant was collected, and the protein concentration was measured using the Pierce™ BCA Protein Assay Kit (Thermo Fisher Scientific). The GFP levels were measured in cell extracts by immunoblotting against GFP.

### Promoter assay by GFP immunoblotting

Equal volumes of protein (obtained using the BCA method) were loaded for SDS-PAGE and immunoblotting. GFPs were detected using the 1-Step™ NBT/BCIP substrate solution (Thermo Fisher Scientific). Signals were recorded using Lumix DMC-GX1 (Panasonic) and quantified using the image analysis tool Fiji (version 2.0.0-rc-65) (48). For quantification, cell lysates of the PpsbA2-GFP strain (KR9) were diluted 2-,4-,8-,16-, and 100-fold with the GT strain and loaded with other cell lysates.

### Antibodies for immunoblotting

For GFP detection, an anti-GFP monoclonal antibody (mFX75 Wako; 1:2,000 dilution) and an alkaline phosphatase-conjugated anti-mouse IgG (ab97020 Abcam; 1:20,000 dilution) were used as primary and secondary antibodies, respectively. To detect the FLAG epitope tag, alkaline-phosphatase-conjugated anti-FLAG IgG (A9469, Sigma Aldrich) was used at a dilution of 1:5,000. For immunoblotting of SigE, a polyclonal antibody against polypeptides from SigE (DGQQGSFNKAVSED) was produced and purified by Eurofins Genomics. Subsequently, the anti-SigE antibody (1:2,000 dilution) was used as the primary antibody, and an alkaline phosphatase-conjugated anti-rabbit IgG antibody (A3937 Sigma Aldrich; 1:20,000 dilution) was used as the secondary antibody. All alkaline-phosphatase-conjugated IgGs were detected using the 1-Step™ NBT/BCIP substrate solution (Thermo Fisher Scientific).

### Genotyping, RNA extraction, and RT-qPCR

Cyanobacterial genomes were extracted using glass beads and vortexed using the phenol-chloroform method (49), and KOD-FXNeo was used for PCR. RNA extraction and RT-qPCR were conducted as previously described (50). The oligonucleotides used for genomic PCR and RT-qPCR are listed in Supplemental Table S2.

### Statistical analysis

Pairwise comparisons were conducted using the Welch’s *t*-test. For multiple comparisons, p values were adjusted using Bonferroni’s method.

### Preparation of the input cell lysate for ChIP

ChIP was conducted using a modified version of a method described in a previous study (51); the names and compositions of buffers are provided in the previous report (51), unless otherwise mentioned. Cells equivalent to those present in 120 mL of culture (eight tubes) were used for the SigE-FLAG ChIP-seq experiment, and cells equivalent to those present in 60 mL of the culture were used for SigA-FLAG ChIP-seq. Each culture of 15 mL was divided into 2-mL tubes along with 100 μL of acid-washed glass beads (150-212 μm; G1145; Sigma-Aldrich) and 50 μL of lysis buffer supplemented with x1 Complete Mini EDTA-free (Roche) and 1 mM PMSF. The cells were bead beaten using the ShakeMaster® NEO for five repetitions of 1 min at 1500 rpm and incubation at 1-min intervals on ice. After bead-beating, 500 μL of lysis buffer was added to two tubes, and the cell lysate was collected. To shear DNA into fragments of 200 bp, the cell lysates were subjected to sonication (24 cycles of 10 s each, 20% amplitude, and incubation at 50-s intervals on ice) performed using VC-750 (EYELA, Tokyo, Japan). After the cell debris was removed by centrifugation (18,000 × *g*, 5 min) at 4°C, the supernatant was collected, and the lysate was diluted in lysis buffer so that total protein concentration was 4 mg /mL.

### ChIP

The names and compositions of buffers are the same as those reported in a previous study, unless otherwise mentioned (51). Prior to immunoprecipitation, an anti-FLAG ® M2 Affinity Gel (A2220, Sigma Aldrich) was equilibrated with PBS supplemented with 3% BSA. For immunoprecipitation, 20 μL of the anti-FLAG ® M2 Affinity Gel was added to the cell lysate during the ChIP-seq assay, followed by rotation for 2 h at 4 °C. The beads were collected by centrifugation (800 × *g*, 1 min) and washed twice with lysis buffer, once with wash buffer 1, twice with wash buffer 2, and once with TE buffer. The beads bound to the DNA-protein complex were eluted using 100 μL of elution buffer (10 mM Tris-HCl (pH 8.0), 100 mM NaCl, and 0.1% SDS) for 30 min at 60 °C. In parallel with immunoprecipitation, 20 μL of the lysate was diluted with 80 μL of the elution buffer and treated as input. The eluted solution and input were treated with 1 mg/mL (final concentration) of RNase A (Nippon Gene, Tokyo, Japan) for 30 min at 37 °C. We added 0.5 mg/mL (final concentration) proteinase K (Kanto Chemical) to the solutions and incubated the solutions overnight at 65 °C. Next, DNA was extracted using the FastGeneM Gel/PCR Extraction Kit (NIPPON Genetics) according to the manufacturer instructions.

### Library preparation and next-generation sequencing

Input and immunoprecipitated DNA were used to prepare multiplexed libraries using the ThruPLEX® DNA-Seq Kit (Takara Bio) and DNA Unique Dual Index Kit (Takara). The multiplexed libraries were dispatched to Eurofins Genomics Inc. and subjected to paired- end sequencing with HiSeqX.

### Mapping and peak calling

The paired-end sequences were mapped onto the *Synechocystis* genome (ASM972v1) using bowtie2 (52)(ver. 2.4.4 paired-end). Peaks were called using the MACS2 program (ver. 2.2.7) with <1e-30 of the q-value cutoff and paired-end option (53), and the intersection of peaks from two replicates excluding the peak region called by control IP was used in subsequent analyses (Fig. S2C). The summits of binding peaks were calculated as the midpoint of the peaks for the two replicates.

### Genome-wide analyses

The positions of the TSS, including internal start sites, were obtained from a previous study (31). The peaks of SigA and SigE are listed in Supplemental Tables S4 and S5, respectively. The intersection and distance of the genomic region were calculated using BEDTools (ver. 2.29.2) (54). We assigned ChIP-seq peaks to TSS within 100 bp of the ChIP-seq peak summit. GO enrichment analysis was performed using a GO search (32, 55) on the gene ontology resource website (http://geneontology.org/). Motif search was performed using the MEME-suite web service (ver. 5.4.1, https://meme-suite.org/meme/). We used the STREME program (33) with the default option for all SigA and SigE peaks, and the MEME program (56) with the default option was used for strong SigA peaks (> 20-fold enrichment), as the number of strong SigA peaks was less than 50.

### List of SigE-dependent genes

The SigE-dependent genes were selected under the following conditions: log_2_ fold change < -0.5 in the *sigE*-deleted strain and > 1.5 in the SigE-overexpressing strain, as indicated in the previous tiling allay data (26, 30). The selected genes were grouped into transcription units (TUs) based on findings reported by a previous study (31).

### Data availability

Raw ChIP sequencing data were deposited in the Sequence Read Archive (accession ID: PRJNA761880).

## Acknowledgements

This study was supported by the following grants to T.O.: Grant-in-Aid for Scientific Research (B) (grant no. 20H02905) and JST-ALCA of the Japan Science and Technology Agency (grant number JPMJAL1306). We thank Dr. Kohki Yoshimoto for providing laboratory instruments, Dr. Shiho Takahasi-Kariyazono for helping preparation of NGS library. We also thank the members of our laboratory. We would like to thank Editage (www.editage.com) for English language editing. Computations were partially performed on the NIG supercomputer at the ROIS National Institute of Genetics.

## Author contributions

RK and TO designed the study; RK performed the experiment; RK analyzed the data; RK and TO wrote the paper.

